# The intraflagellar transport protein IFT20 recruits ATG16L1 to early endosomes to promote autophagosome formation in T cells

**DOI:** 10.1101/2020.11.26.400135

**Authors:** Francesca Finetti, Chiara Cassioli, Valentina Cianfanelli, Fabrizia Zevolini, Anna Onnis, Monica Gesualdo, Jlenia Brunetti, Francesco Cecconi, Cosima T. Baldari

## Abstract

Lymphocyte homeostasis, activation and differentiation crucially rely on basal autophagy. The fine-tuning of this process depends on autophagy-related (ATG) proteins and their interaction with the trafficking machinery that orchestrates the membrane rearrangements leading to autophagosome biogenesis. The underlying mechanisms are as yet not fully understood. The intraflagellar transport (IFT) system, known for its role in cargo transport along the axonemal microtubules of the primary cilium, has emerged as a regulator of autophagy in ciliated cells. Growing evidence indicates that ciliogenesis proteins participate in cilia-independent processes, including autophagy, in the non-ciliated T cell. Here we investigate the mechanism by which IFT20, an integral component of the IFT system, regulates basal T cell autophagy. We show that IFT20 interacts with the core autophagy protein ATG16L1 through its CC domain, which is also essential for its pro-autophagic activity. We demonstrate that IFT20 is required for the association of ATG16L1 with the Golgi complex and early endosomes, both of which have been identified as membrane sources for phagophore elongation. This involves the ability of IFT20 to interact with proteins that are resident at these subcellular localizations, namely the golgin GMAP210 at the Golgi apparatus and Rab5 at early endosomes. GMAP210 depletion, while leading to a dispersion of ATG16L1 from the Golgi, did not affect basal autophagy. Conversely, IFT20 was found to recruit ATG16L1 to early endosomes tagged for autophagosome formation by the BECLIN 1/VPS34/Rab5 complex, which resulted in the local accumulation of LC3. Hence IFT20 participates in autophagosome biogenesis under basal conditions by regulating the localization of ATG16L1 at early endosomes to promote autophagosome biogenesis. These data identify IFT20 as a new regulator of an early step of basal autophagy in T cells.

## INTRODUCTION

Autophagy is a degradative process that subserves the dual function of eliminating damaged macromolecules and organelles while providing endogenous energy sources and building blocks to maintain cellular homeostasis (Dikic and Elazar, 2018). Similar to other cell types, T cells exploit autophagy as a quality control and pro-survival mechanism which is essential for the long-lasting maintenance of the naive peripheral T cell repertoire. The roles of autophagy are however not limited to T cell homeostasis. Autophagy participates in thymocyte development by sustaining the survival of double negative thymocytes and their transition to the double positive stage (Nedjic et al., 2008) and regulates both positive and negative selection via the MHCII loading pathway in thymic epithelial cells (Kasai et al., 2009). In the periphery, T cell activation and proliferation are fine-tuned by selective autophagy (Zaffagnini and Martens, 2016). In this context, the autophagy cargo is initially switched from mitochondria to macromolecules including inhibitors of T cell receptor (TCR) signaling (Hubbard et al., 2010; Valdor et al., 2014) and cell cycle (Jia et al., 2015). Subsequently, as T cells undergo differentiation, the autophagic degradation of the NF-κB regulator Bcl10 occurs (Paul et al., 2012). Autophagy also impacts on helper and cytotoxic T cell effectors, with Th1, Th2 and Treg cells relying on autophagy for survival (Kabat et al., 2016; Kovacs et al., 2012; Wei et al., 2016) and cytotoxic T lymphocytes (CTL) for their function and for memory maintenance (Puleston et al., 2014; Xu et al., 2014).

The autophagy machinery has been extensively characterized. Autophagy is initiated by the ULK complex, consisting of ULK1/2, ATG13, FIP200 and ATG101, which promotes phagophore nucleation through the class III PI3-K complex, consisting of VPS34, VPS15, BECLIN 1 and ATG14L. In order to engulf the cargo, the phagophore elongates and subsequently closes. These two steps involve the ATG12-ATG5 conjugation system, which is recruited to the phagophore by ATG16L. The conjugation system acts as an E3-like ligase to promote the cleavage and lipidation of ATG8/LC3, resulting in the phosphatidyl ethanolamine-conjugated form that binds to the phagophore membrane. Following phagophore closure, the resulting autophagosomes undergo dynein-dependent transport along the microtubules to a perinuclear location, where they fuse with lysosomes to become autolysosomes, delivering their cargo for degradation (Dikic and Elazar, 2018). This basic process is fine-tuned by a plethora of regulators and effectors. Among these is the intraflagellar transport (IFT) system, a multimolecular protein complex which controls the assembly of the primary cilium, a signaling organelle present in the majority of vertebrate cells (Pampliega et al., 2013; Prevo et al., 2017). Two integral components of the IFT system, IFT20 and IFT88, have been shown to mediate the transport of some components of the autophagic machinery to the cilium during cell starvation, with IFT20 binding ATG16L1 at the Golgi apparatus and shuttling it to the base of the cilium, wherefrom it enters the cilium with the assistance of IFT88 (Pampliega et al., 2013).

We have previously shown that, unexpectedly, IFT20 plays a key role in the activation of the non-ciliated T cells in a complex with other IFT components by regulating the assembly of the immune synapse, a specialized signaling interface that forms when a T cell encounters a cognate antigen presenting cell (Finetti et al., 2009, 2011). This function involves the ability of IFT20 to associate with the Golgi complex and endocytic compartments, the primary one among the latter being the early endosome, to control the intracellular traffic of the T cell antigen receptor and the transmembrane adaptor linker for activation of T cells (LAT) (Finetti et al., 2009, 2014; Vivar et al., 2016). Interestingly, we have recently implicated IFT20 in another vesicular trafficking-related process, the mannose-6-phosphate receptor-dependent transport of acid hydrolases to lysosomes, on which lysosome biogenesis and function depend (Finetti et al., 2020). Consistent with the central role of lysosomes in autophagy, both basal and starvation-induced autophagy are impaired in T cells (Finetti et al., 2020). However, the implication of the IFT system at early steps of autophagy in ciliated cells (Pampliega et al., 2013) suggests that IFT20 could participate in T cell autophagy also directly. Here we show that IFT20 promotes ATG16L1 localization to the Golgi complex and early endosomes through its interaction with GMAP210 and Rab5, respectively. We demonstrate that under basal conditions IFT20 participates in autophagosome formation at early endosomes, but not at the Golgi apparatus, by allowing for the recruitment of ATG16L1 to early endosome-associated BECLIN 1 complex, thereby promoting local LC3-II accumulation. Our results identify IFT20 as an adaptor that targets ATG16L1 and downstream autophagy regulators to a specific endomembrane compartment, resulting in autophagosome biogenesis in T cells.

## MATERIALS AND METHODS

### Cells, plasmids, and transfections

Control, IFT20KD and GMAPKD Jurkat T cell lines were generated as previously described (Finetti et al., 2009; Galgano et al., 2017). Jurkat T cells were stably transfected with the pEGFP-N1 plasmid construct encoding full-length IFT20, or a deletion mutant of IFT20 lacking aminoacid residues 73-132, which include the coiled-coil domain (ΔCC IFT20), or the respective empty vector. Stably transfected cells were selected in G418-containing medium at the final concentration of 1 mg/ml (Gibco-BRL/Life Technologies). Transient transfections were carried out by electroporation using pCMV-EGFP-C3-Rab5a (kindly provided by M. Zerial), pEGFP 2xFYVE (kindly provided by A. De Matteis), pEGFP-N1 IFT20-GFP or pEGFP-N1 ΔCC IFT20-GFP and analysed 24 h post-transfection.

### Cloning and purification of recombinant proteins

GFP- and GST-tagged mutants of IFT20 were generated by cloning the sequences that encode the IFT20 N-terminus lacking the coiled-coil domain (ΔCC-IFT20, aa 1-73), the IFT20 C-terminus including the coiled-coil domain (CC-IFT20, aa 74-132), and the full-length protein (IFT20, aa 1-132) in-frame with the tags into the pEGFP-N1 (#6085-1 Addgene) and pGEX-6P-2 vectors (#27-4598-01Addgene). The sequences were amplified by PCR using the primers listed in Supplementary Table 1. The 5’-ends of the primers were modified to add compatible restriction sites (XhoI and KpnI for pEGFP-N1, EcoRI and XhoI for pGEX-6P-2) and extra base pairs that ensure efficient DNA cleavage by restriction enzymes. The recombinant GST fusion proteins were affinity purified on GSH-Sepharose (GE Healthcare) from bacterial cultures incubated with 0.25 mM isopropyl-β-D-thiogalactopyranoside (Sigma-Aldrich) overnight at RT and lysed in B-PER Bacterial Protein Extraction Reagent (Thermo Fisher Scientific).

### Antibodies and reagents

All primary commercial antibodies used in this manuscript are listed in Table S2, where information about the dilutions used for immunoblotting and immunofluorescence is specified. Polyclonal anti-IFT20 antibodies (Pazour et al., 2002) were kindly provided by G. Pazour. Secondary peroxidase-labeled antibodies were from Amersham Biosciences. Alexa Fluor 488- and 555-labeled secondary Abs were from ThermoFisher Scientific (anti-mouse 488, #A11001; anti-rabbit 488, #A11008; anti-mouse 555, #A21422; anti-rabbit 555, #A21428). Chloroquine was purchased from Sigma-Aldrich (C6628).

### Immunoprecipitation, *in vitro* binding assays and Immunoblotting

Immunoprecipitation experiments were performed as previously described (Finetti et al., 2020). Briefly, 5×10^7^ cells/sample were lysed in 0.5% Triton X-100 in 20 mM Tris-HCl (pH 8), 150 mM NaCl in the presence of protease inhibitors (Sigma-Aldrich) and the phosphatase inhibitor sodium vanadate (Sigma-Aldrich). Postnuclear supernatants (2 mg/sample) were immunoprecipitated for 2 h using 2 μg of rabbit anti-IFT20 antibody (#13615-1-AP, Proteintech), anti-ATG16L1 antibody (#8089S, Cell Signaling) or mouse anti-BECLIN 1 mAb (sc-48341, Santa Cruz), and protein A-Sepharose (PAS, 3 mg/sample, GE Healthcare), after a preclearing step on PAS (1 h, 3 mg/sample). Subsequently, all samples were washed 4X with 1 ml 0.5% Triton X-100 lysis buffer, resuspended in 15 μl Laemmli buffer (#B0007, Life Technologies), boiled for 5 min and then subjected to SDS-PAGE.

In vitro-binding assays were carried out using recombinant GST, IFT20-GST, ΔCC IFT20-GST or CC IFT20-GST on GSH-Sepharose precleared postnuclear supernatants from 5×10^7^ cells/sample lysed in 0.5% Triton X-100 in the presence of protease and phosphatase inhibitors as described (Pacini et al., 1998).

Immunoblotting was carried out using peroxidase-labeled secondary antibodies and a chemiluminescence detection kit (#34078, Pierce Rockford). Membranes were reprobed with control antibodies after stripping carried out using ReBlot Plus Mild Antibody Stripping Solution, 10x (#2502, Merck Millipore). Blots were scanned using a laser densitometer (Duoscan T2500; Agfa) and quantified by using ImageJ 1.46r (National Institutes of Health, USA).

### Immunofluorescence microscopy and colocalization analyses

For immunofluorescence analysis, Jurkat cells were allowed to adhere for 15 min to poly-L-lysine-coated wells and then fixed by immersion in methanol for 10 min at −20°C. Alternatively, for LC3B labelling, Jurkat cells were incubated 10 min in 50 mM NH4Cl after fixation in 4% paraformaldehyde for 10 min at RT, and then permeabilized in methanol for 10 min at −20°C. Fixed and permeabilized samples were washed for 5 min in PBS and incubated with primary antibodies overnight at 4°C, after blocking in 5% normal goat serum and 1% BSA for 1 h. Samples were washed for 5 min in PBS and incubated for 45 min at RT with Alexa-Fluor-488-, Alexa-Fluor-555- and Alexa-Fluor-647-labeled secondary antibodies.

Confocal microscopy was carried out on a Zeiss LSM700 or a TCS SP8 Confocal laser scanning microscopy (Leica) using a 63X objective with pinholes opened to obtain 0.8 μm-thick sections. Detectors were set to detect an optimal signal below the saturation limits. Images were processed with Zen 2009 image software (Carl Zeiss, Jena, Germany). The quantitative colocalization analysis was performed on median optical sections using ImageJ and JACoP plug-in to determine Mander’s coefficient as previously described (Finetti et al., 2014). The ATG16L1 dispersion was quantified by measuring fluorescence intensity in concentric regions using ImageJ. Circular regions were centered on the point of ATG16L1 maximal intensity and designed proportionally to the cell size (inner circle, middle ring and outer ring diameters, corresponding to 1/9, 1/4.5 and 1/2.25 of cell diameter, respectively) (Progida et al., 2010).

### Autophagic flux and LC3^+^ dot number measurement

To analyse the autophagic flux, 1×10^6^ cells/sample were incubated in RPMI 1640 added with 7.5% BCS for 30 minutes at 37°C in the presence or absence of 40 μM chloroquine. Subsequently, cells were harvested and lysed in 1% Triton X-100 in 20 mM Tris-HCl pH 8.0, 150 mM NaCl in the presence of protease and phosphatase inhibitors, and processed for immunoblotting with anti-LC3B antibodies. Autophagy flux was calculated as the difference in LC3-II levels, normalized to actin, between chloroquine-treated and untreated cells. The number of LC3B^+^ vesicles was determined by immunofluorescence microscopy. 1×10^5^ cells/sample were incubated in RPMI 1640 added with 7.5% BCS for 30 minutes at 37°C with or without 40 μM chloroquine and allowed to adhere to poly-L-lysine-coated wells. Subsequently, the samples were fixed, permeabilized and stained as described above.

### Membrane fractionation

Cytosolic and membrane fractions were purified as previously described (Finetti et al., 2014). Jurkat cells (3×10^7^/sample) were resuspended in 1 ml homogenization medium (0.25 M sucrose, 1 mM EDTA, 10 mM Tris-HCl pH 7.4) in the presence of protease and phosphatase inhibitors. The samples were homogenized by 10 pestle strokes through Dounce homogenization (tight Dounce homogenizer, Wheaton, USA) and 10 passages through a 26-gauge syringe needle. The homogenate was centrifuged at 3,000 g for 5 min at 4°C to remove nuclei and the supernatant was centrifuged at 65,000 g for 1 h at 4°C. The supernatant (cytosolic fraction) was collected, while the pellet (membrane fraction) was lysed in homogenization buffer containing protease and phosphatase inhibitors with 0.5% Triton, and centrifuged at 16,100 g for 20 min at 4°C to eliminate insoluble material. The same quantities of membrane protein-enriched supernatant and cytosolic fraction were analysed by SDS-PAGE.

### Statistical analysis

GraphPad (Prism Software) was used to calculate mean values, standard deviation values and statistical significance. Values with Gaussian distribution were analysed using Student’s t test (paired or unpaired), one sample t test (theoretical mean=1) or one-way ANOVA. Values without normal distribution were analysed using Mann-Withney test or Kruskal-Wallis test. A level of P<0.05 was considered statistically significant.

## RESULTS

### IFT20 interacts constitutively with ATG16L1 in T cells

IFT20 interacts with ATG16L1 and colocalizes with this protein at intracellular vesicles in ciliated cells (Pampliega et al., 2013). The evidence that T cells share common regulators of vesicular trafficking with ciliated cells (Cassioli and Baldari, 2019) raises the question of whether IFT20 may interact with ATG16L1 also in non-ciliated T cells. To address this issue, we performed co-immunoprecipitation experiments in Jurkat T cells, which showed that IFT20 interacts with ATG16L1 (Figure 1A). Consistent with this finding, IFT20 displayed partial colocalization with ATG16L1 in Jurkat T cells, as assessed by immunofluorescence (Figure 1B). The ability of IFT20 to interact with ATG16L1 was confirmed by *in vitro* GSH-Sepharose pull-down assays using an IFT20-GST fusion protein (Figure 1C,D). Hence, similar to ciliated cells, IFT20 interacts with ATG16L1 in the non-ciliated T cell.

IFT20 is a small protein containing a single coiled-coil (CC) domain at its C-terminus (Figure 1C) that has been implicated in its heterodimerization with other CC domain-containing components of the IFT complex (Baker et al., 2003; Omori et al., 2008). To map the interaction of ATG16L1 on IFT20 we generated GST fusion proteins with the IFT20 CC domain (aminoacid residues 74-132; CC IFT20) or the full-length protein lacking the CC domain (aminoacid residues 1-73; ΔCC IFT20). The GST-pull down assays showed that ATG16L1 binds IFT20 regardless of the presence of the CC-domain (Figure 1D). These results indicate that IFT20 interacts with ATG16L1 through a molecular determinant localized within its unstructured region, potentially using its CC domain to couple ATG16L1 to sites of autophagosome formation.

**Figure 1.**
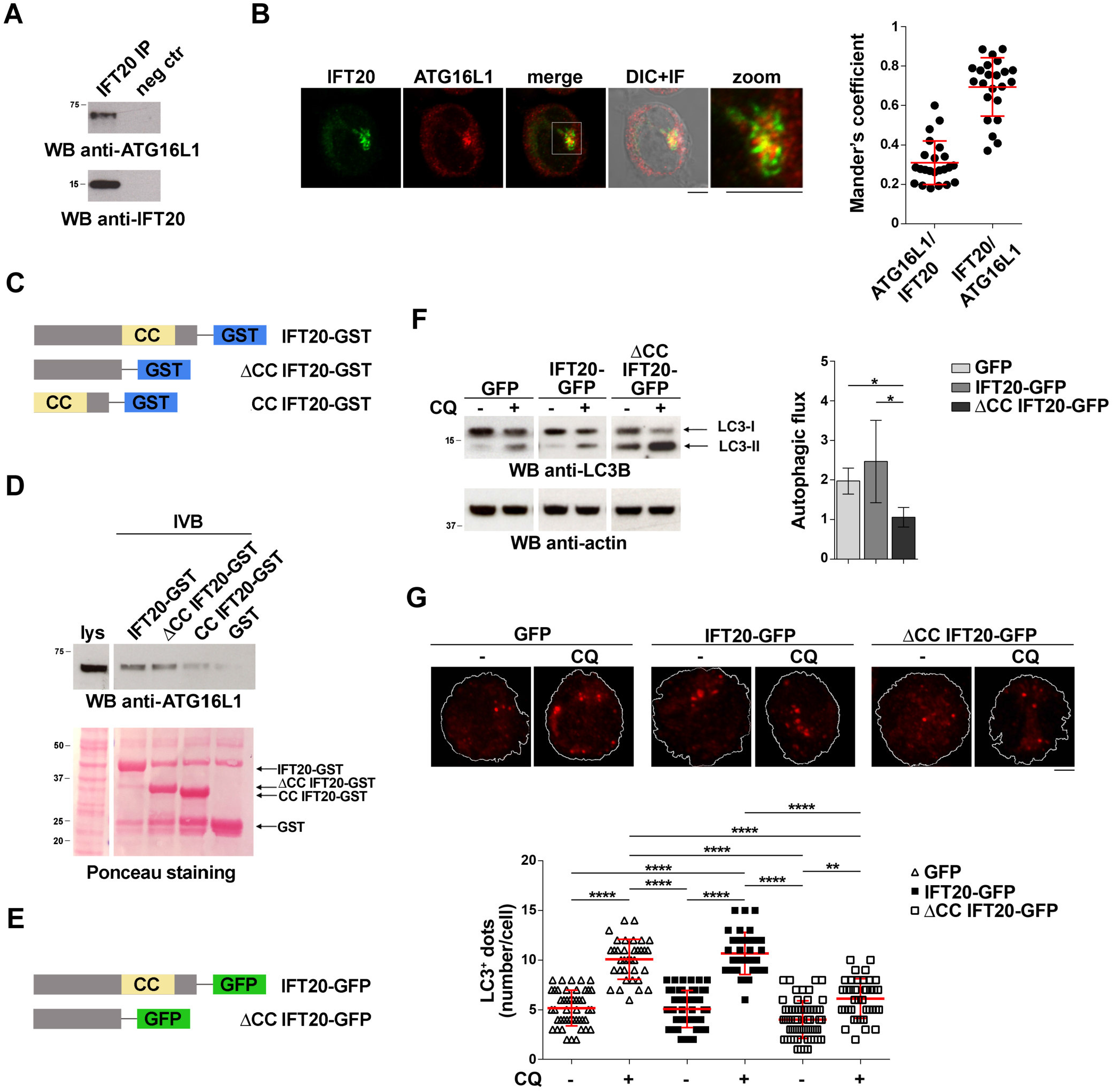
IFT20 binds to ATG16L1 and promotes T cell autophagy through its CC domain. (A) Western blot (WB) analysis with anti-ATG16L1 antibodies of IFT20-specific immunoprecipitates from lysates of Jurkat T cells. A preclearing control (proteins bound to Protein-A–Sepharose before the addition of primary antibody) is included in each blot (neg ctr). The migration of molecular mass markers is indicated. The immunoblots shown are representative of at least 3 independent experiments. (B) Immunofluorescence analysis of IFT20 and ATG16L1 in Jurkat cells. Representative medial optical sections and overlay of immunofluorescence (IF) and differential interference contrast (DIC) images are shown (IF+DIC). The graph shows the quantification (using Mander’s coefficient) of the weighted colocalization of IFT20 and ATG16L1 in Jurkat cells. The data are expressed as mean±s.d. (23 cells/sample; n=3). Scale bars: 5 μm. (C) Schematic representation of the GST fusion proteins with full-length IFT20 (IFT20-GST), or the full-length protein lacking the CC domain (ΔCC IFT20-GST) or the CC domain (CC IFT20-GST). The CC domain is highlighted as a yellow box. (D) Immunoblot analysis with anti-ATG16L1 antibodies of in vitro-binding assays carried out on post-nuclear supernatants of Jurkat cells using IFT20-GST, ΔCC IFT20-GST and CC IFT20-GST fusion proteins, or GST as negative control. The Ponceau red staining of the same filter is shown to compare the levels of fusion proteins and GST used in the assay. The immunoblot shown is representative of three independent experiments. (E) Schematic representation of the GFP fusion protein with IFT20 (IFT20-GFP) or with the full-length protein lacking the CC domain (ΔCC IFT20-GFP). The CC domain is highlighted as a yellow box. (F) Immunoblot analysis of LC3B in lysates of Jurkat cells expressing GFP, IFT20-GFP or ΔCC IFT20-GFP in the presence or absence of chloroquine (CQ, 40 μM). The migration of molecular mass markers is indicated. The graph shows the autophagic flux in Jurkat transfectants, calculated as the difference in the levels of LC3II/actin between CQ-treated and CQ-untreated samples (mean fold ± SD; one-way ANOVA; n ≥ 3). (G) Immunofluorescence analysis of LC3B in Jurkat cells expressing GFP, IFT20-GFP or ΔCC IFT20-GFP either untreated or treated for 30 min with chloroquine (CQ, 40 μM). The z-projection of maximum intensity and the quantification of the number of LC3^+^ dots/cell are shown. At least 35 cells from three independent experiments were analysed (mean ± SD; Kruskal–Wallis test). *P < 0.05; **P < 0.01; ****P < 0.0001.

To investigate the role of the CC domain in basal IFT20-dependent autophagy we generated Jurkat T cells transfectants expressing IFT20-GFP or a deletion mutant lacking the CC domain, fused to GFP (ΔCC IFT20-GFP) (Figure 1E and Supplementary figure1A). A similar construct encoding the isolated GFP-tagged CC domain was not expressed at detectable levels, likely due to low protein stability (data not shown). The impact of ΔCC-IFT20 expression on the autophagic flux was assessed by immunoblot. Cells were either untreated or treated with chloroquine, which blocks the degradation of LC3-II by impairing autophagosome fusion with lysosomes (Mauthe et al., 2018). Immunoblot analysis with anti-LC3 antibodies showed that under basal conditions the autophagic flux was decreased in cells expressing ΔCC IFT20-GFP compared to controls expressing full-length IFT20-GFP (Figure 1F). This result was confirmed by immunofluorescence analysis of LC3^+^ dots, which showed a decreased autophagic flux in ΔCC IFT20-GFP-expressing cells compared to controls expressing the full-length GFP fusion protein (Figure 1G). Hence, the pro-autophagic function of IFT20 requires its CC domain.

### IFT20 tethers ATG16L1 to cellular membranes

ATG16L1 is a cytoplasmic protein that is recruited to multiple cellular membranes which can serve as membrane sources for autophagosomes (Xiong et al., 2018). Although the mechanism of ATG16L1 recruitment to membranes is not fully understood, ATG16L1 has been reported to interact with the autophagy regulator FIP200 and with the phosphatidylinositol 3-phosphate (PI(3)P)-binding protein WIPI2. Both these interactions are functional to the association of ATG16L1 with isolation membranes, particularly at the endoplasmic reticulum, during autophagosome formation. ATG16L1 also interacts with the small GTPase Rab33 at the Golgi complex, SNX18/Rab11 at recycling endosomes, and the membrane coat protein clathrin at the plasma membrane, wherefrom it is transferred to the endocytic compartment (Gammoh et al., 2013; Itoh et al., 2008; Ravikumar et al., 2010).

The association of IFT20 with the Golgi apparatus and endocytic compartments (Finetti et al., 2009) and its ability to interact with ATG16L1 (Figure 1) suggest a role for IFT20 in tethering ATG16L1 to endomembranes. To test this hypothesis we addressed the outcome of RNAi-mediated IFT20 depletion (IFT20KD; Supplementary figure 1B) on the subcellular localization of ATG16L1. Immunofluorescence analysis showed that ATG16L1 was concentrated at a vesicular-like compartment in control cells, as previously reported in NIH3T3 cells (Itoh et al., 2008). The localization of ATG16L1 was altered to a significant extent in Jurkat cells stably knocked down for IFT20 expression, where it showed a dispersed pattern, as assessed by quantifying the fluorescence intensity in concentric cellular regions from the point of maximal intensity (Figure 2A) (Progida et al., 2010). This result supports the notion that IFT20 contributes to the vesicular localization of ATG16L1 in T cells.

**Figure 2.**
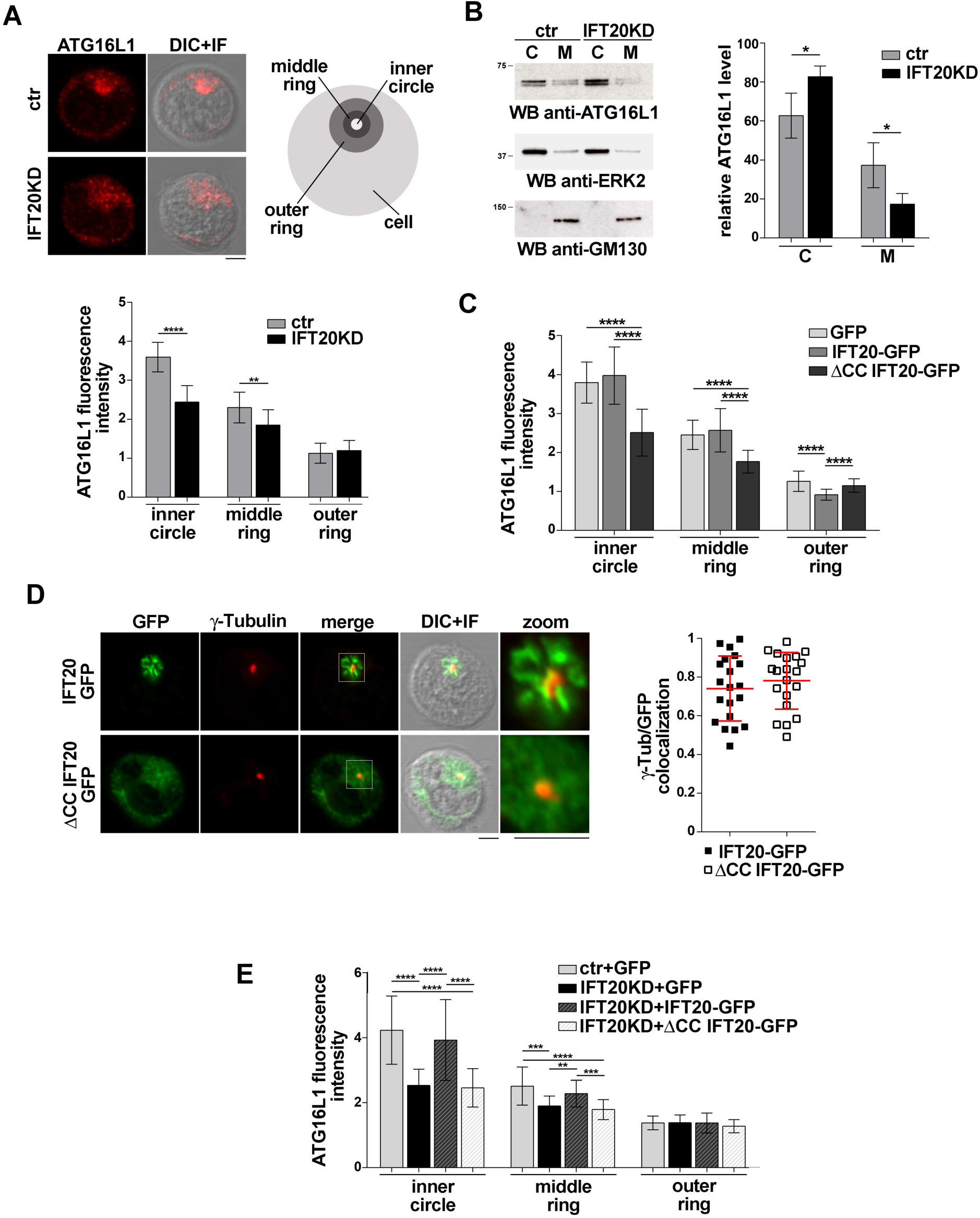
IFT20 regulates the intracellular localization of ATG16L1. (A) Immunofluorescence analysis of ATG16L1 in control (ctr) and IFT20KD Jurkat cells. Representative medial optical sections and overlay of immunofluorescence and DIC images are shown (IF+DIC). The histogram below shows the quantification of fluorescence intensity in the concentric regions indicated in the scheme and defined from the point of ATG16L1 maximal intensity (mean±s.d, ≥20 cells/sample; n=3; Mann-Whitney test). Scale bars: 5 μm. (B) Immunoblot analysis of ATG16L1 in cytosolic (C) and membrane (M) fractions purified from control and IFT20KD Jurkat cells. The cytosolic protein ERK2 and the cis-Golgi marker GM130 were used to assess the purity of cytosolic and membrane fractions, respectively. The migration of molecular mass markers is indicated. The histogram shows the quantification of the percentage of ATG16L1 in the cellular fractions obtained from 4 independent experiments (mean±s.d; Student’s t-test). (C) Immunofluorescence analysis of ATG16L1 in GFP, IFT20-GFP and ΔCC IFT20-GFP Jurkat transfectants. The histograms show the quantification of fluorescence intensity in the concentric regions described above (mean±s.d, ≥25 cells/sample; n=3; Mann–Whitney test). (D) Quantification (using Mander’s coefficient) of the weighted colocalization of γ-tubulin with GFP in medial confocal sections of IFT20-GFP or ΔCC IFT20-GFP expressing Jurkat cells (mean±s.d.; ≥20 cells/line; n=3). Representative images (medial optical sections and overlay DIC+IF) are shown. Scale bar: 5 μm. (E) Immunofluorescence analysis of ATG16L1 in control and IFT20KD cells transiently transfected with either empty vector (GFP), or the IFT20-GFP construct or the ΔCC IFT20-GFP construct. The graph shows the quantification of fluorescence intensity in the concentric regions described above (mean±s.d, ≥25 cells/sample; n=3; Mann–Whitney test). *P < 0.05; **P < 0.01; ***P < 0.001; ****P < 0.0001

To confirm this function of IFT20 the association of ATG16L1 with cellular membranes was addressed biochemically by immunoblot analysis of cytosolic and membrane fractions obtained from control and IFT20KD Jurkat cells. ATG16L1 was detected both in cytosolic and in membrane fractions. IFT20 deficiency resulted in a decrease in the amount of ATG16L1 associated with the membrane fractions (Figure 2B), suggesting that the loss of ATG16L1 compartmentalization observed by immunofluorescence in IFT20KD cells results from its release from cellular membranes to the cytosol.

To understand whether the CC domain of IFT20, which is required for the pro-autophagic function of IFT20 (Figure 1F,G) but not for ATG16L1 binding (Figure 1D), affects the subcellular localization of ATG16L1, we carried out an immunofluorescence analysis of ATG16L1 in Jurkat T cells expressing IFT20-GFP or the ΔCC IFT20-GFP mutant (Figure 1E and Supplementary figure 1A). At variance with IFT20-GFP, which did not elicit any detectable effect, the absence of the CC domain resulted in a complete loss of the vesicular pattern of ATG16L1 (Figure 2C and 4E), similar to that observed in IFT20KD cells (Figure 2A). The alteration in the subcellular localization of ATG16L1 was paralleled by a diffuse staining throughout the cell of ΔCC IFT20-GFP, with only the centrosomal localization remaining unaffected (Figure 2D). This suggests that ΔCC IFT20-GFP acts in a dominant negative fashion to prevent the interaction of endogenous, membrane-associated IFT20 with ATG16L1. Importantly, the defect in ATG16L localization observed in IFT20KD cells was rescued by restoring IFT20 expression, while ΔCC-IFT20 was unable to correct this defect (Figure 2E and Supplementary figure 1C). Hence IFT20 mediates the vesicular localization of ATG16L in T cells and this function maps to its CC domain.

### IFT20 is required for the association of ATG16L1 with the Golgi apparatus

To identify the membrane compartment ATG16L1 associates with in T cells, we carried out a colocalization analysis of ATG16L1 with the Golgi complex, one of the most IFT20-enriched subcellular compartments (Finetti et al., 2009; Follit et al., 2006). A decrease in the colocalization of ATG16L1 with the Golgi marker giantin was observed in IFT20KD cells compared to controls (Figure 3A), implicating IFT20 in the Golgi localization of ATG16L1.

**Figure 3.**
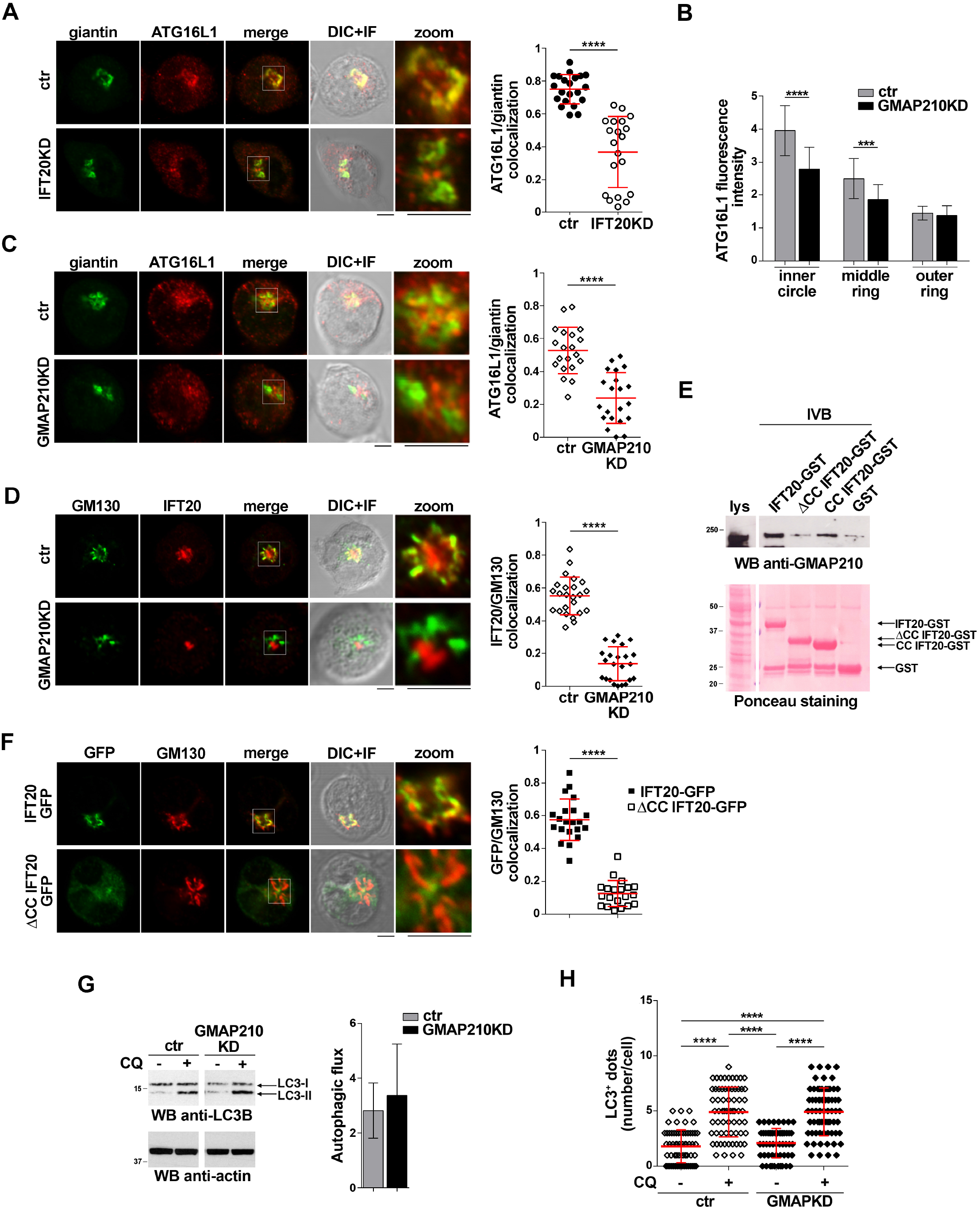
IFT20 couples ATG16L1 to the Golgi through its CC domain-mediated interaction with GMAP210. (A) Quantification using Mander’s coefficient of the weighted colocalization of ATG16L1 and the Golgi marker giantin in ctr and IFT20KD Jurkat cells (≥20 cells/sample, n=3; mean±s.d; Mann–Whitney test). Representative images (medial optical sections and the overlay DIC+IF) are shown. Scale bar: 5 μm. (B) Quantification of fluorescence intensity of ATG16L1 in the concentric regions previously described (Figure 2 A) in control and GMAP210KD cells (mean±s.d, ≥25 cells/sample; n=3; Mann–Whitney test). (C,D) Immunofluorescence analysis of ATG16L1 and giantin (C) or IFT20 and the Golgi marker GM130 (D) in control and GMAP210KD cells. Representative medial optical sections and overlay of immunofluorescence (IF) and differential interference contrast (DIC) images are shown (IF+DIC). The graph shows the quantification (using Mander’s coefficient) of the weighted colocalization of ATG16L1 and giantin (C) or IFT20 and GM130 (D). The data are expressed as mean±s.d. (≥20 cells/sample; n=3; Mann–Whitney test). Scale bars: 5 μm. (E) Immunoblot analysis with anti-GMAP210 antibodies of in vitro-binding assays carried out on post-nuclear supernatants of Jurkat cells using IFT20-GST, ΔCC IFT20-GST and CC IFT20-GST fusion proteins, or GST as negative control. The Ponceau red staining of the same filter is shown to compare the levels of fusion proteins and GST used in the assay. The immunoblot shown is representative of three independent experiments. (F) Quantification (using Mander’s coefficient) of the weighted colocalization of the cis-Golgi marker GM130 with GFP in medial confocal sections of IFT20-GFP or ΔCC IFT20-GFP expressing Jurkat cells (mean±s.d.; ≥20 cells/line; n=3; Mann–Whitney test). Representative images (medial optical sections and overlay DIC+IF) are shown. Scale bar: 5 μm. (G) Immunoblot analysis of LC3B in lysates of control or GMAP210KD cells in the presence or absence of chloroquine (CQ, 40 μM). The migration of molecular mass markers is indicated. The histograms show the autophagic flux calculated as the difference in the levels of LC3II/actin between CQ-treated and CQ-untreated samples (mean fold ± SD; Student’s t-test; n ≥ 3). (H) Quantification of the number of LC3^+^ dots/cell in control or GMAP210KD cells either untreated or treated for 30 min with chloroquine (CQ, 40 μM). At least 35 cells from three independent experiments were analysed (mean ± SD; Kruskal–Wallis test). **P < 0.01; ***P < 0.001; ****P < 0.0001.

IFT20 is known to associate with the Golgi apparatus through its interaction with the golgin GMAP210 (Follit et al., 2008). This interaction is mediated by the CC domain of GMAP210 (Follit et al., 2008; Zucchetti et al., 2019). To understand whether the IFT20-dependent association of ATG16L with the Golgi is mediated by GMAP210 we analyzed the subcellular localization of ATG16L in Jurkat cells knocked down for GMAP210 expression by RNA interference (Supplementary figure 1D). Similar to IFT20KD cells, GMAP210KD cells showed a dispersed pattern of ATG16L1 staining, with a decrease in its association with the Golgi apparatus (Figure 3B, C). This was paralleled by a loss of IFT20 co-localization with the Golgi (Figure 3D), consistent with the ability of IFT20 to interact with GMAP210 in T cells (Galgano et al., 2017; Zucchetti et al., 2019).

To map the GMAP210 binding site on IFT20 we carried out pull-down assays using the IFT20-GST, IFT20-ΔCC-GST and IFT20-CC-GST fusion proteins (Figure 1C). As expected, GMAP210 was found to interact with full-length IFT20 (Figure 3E). The CC domain of IFT20 was largely responsible for this interaction, as shown by the preferential binding to GMAP210 to the isolated CC domain compared to the fusion protein lacking the CC domain (Figure 3E). Consistent with this finding, the CC domain of IFT20 was essential for its association with the Golgi apparatus, as assessed in Jurkat cell transfectants expressing ΔCC IFT20-GFP (Figure 3F).

The Golgi apparatus has been identified as a source of autophagosomal membranes, contributing to phagophore elongation through ATG16L recruitment (Itoh et al., 2008; Staiano and Zappa, 2019). To understand whether the GMAP210-dependent, IFT20-mediated recruitment of ATG16L1 to the Golgi is implicated to the pro-autophagic function of IFT20 we measured the autophagic flux in GMAP210KD cells. Remarkably, GMAP210 deficiency did not impair basal autophagy, as assessed both by immunoblot and by immunofluorescence analysis of LC3 (Figure 3G and 3H). Hence, while IFT20 couples ATG16L1 to the Golgi in T cells through its CC domain-dependent interaction with GMAP210, the resulting Golgi association of ATG16L does not lead to autophagosome formation. Accordingly, the recruitment of BECLIN 1 to the Golgi complex is minimal in Jurkat cells, as assessed by immunofluorescence analysis of the colocalization of BECLIN 1 with the cis-Golgi marker GM130 (Supplementary figure 2).

### IFT20 is required for the association of ATG16L1 with early endosomes

In T cells IFT20 is associated not only with the Golgi apparatus, but also with other endomembrane compartments, a major one being early endosomes, where it interacts with Rab5 (Finetti et al., 2014). Of relevance, Rab5 is a component of the macromolecular complex containing BECLIN 1 and the class III phosphatidylinositol-3 kinase VPS34 that regulates autophagosome formation (Ravikumar et al., 2008). An association of ATG16L1 with both early and recycling endosomes has been reported in other cell types (Fraser et al., 2019; Puri et al., 2013; Ravikumar et al., 2010), suggesting the possibility that IFT20 may exploit its interaction with Rab5 to recruit ATG16L1 to early endosomes to promote autophagy.

To investigate whether ATG16L1 associates with early endosomes in T cells and whether IFT20 is required for its localization, we carried out a colocalization analysis of ATG16L1 and Rab5 in control and IFT20KD T cells. Interestingly, we found that a pool of ATG16L1 was associated with Rab5^+^ endosomes in control cells, as reported for glial cells (Fraser et al., 2019). IFT20 depletion resulted in a significant decrease in the colocalization of ATG16L1 with Rab5 (Figure 4A). Additionally, ATG16L1 was found to interact with Rab5 in co-immunoprecipitation experiments (Figure 4B). This interaction was impaired in IFT20 KD cells (Figure 4B).

**Figure 4.**
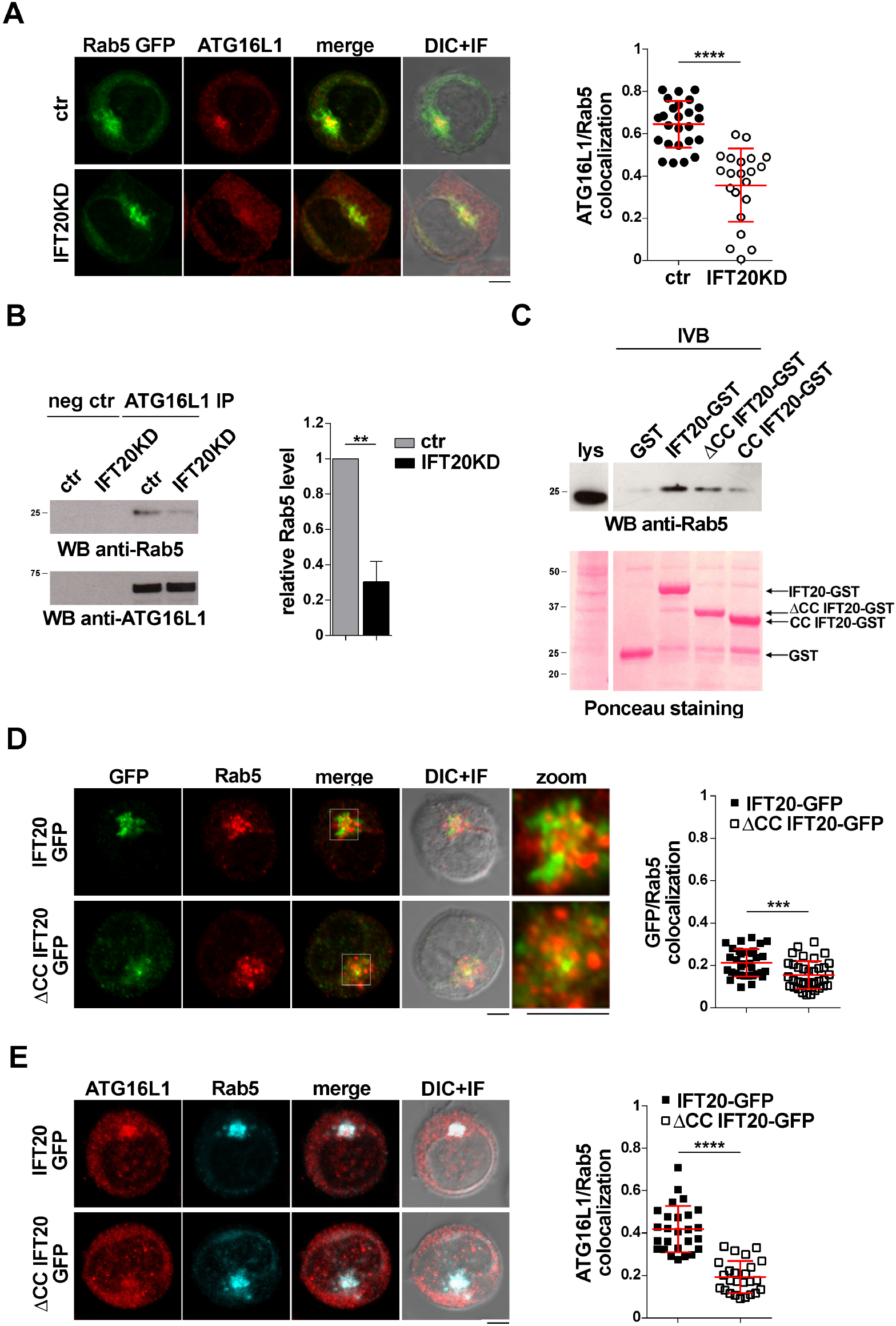
IFT20 couples ATG16L1 to early endosomes through interaction with Rab5. (A) Immunofluorescence analysis of ATG16L1 and Rab5 GFP in control and IFT20KD Jurkat cells. Representative medial optical sections and overlay of immunofluorescence and DIC images are shown (IF+DIC). Scale bar: 5 μm. The graph shows the quantification of the weighted colocalization (Mander’s coefficient) of ATG16L1 and the early endosome marker Rab5 in ctr and IFT20KD Jurkat cells (≥21 cells/sample, n=3; mean±s.d; Mann–Whitney test). (B) Immunoblot analysis with anti-Rab5 antibodies of ATG16L1-specific immunoprecipitates from lysates of control and IFT20KD Jurkat cells. Preclearing controls are included in each blot (neg ctr). Tested proteins show comparable expression in total cell lysate from ctr and IFT20KD Jurkat cells (Supplementary figure 3A and B). The migration of molecular mass markers is indicated. The quantification of the relative protein expression normalized to ATG16L1 (mean fold ± SD; ctr value=1) is reported for each blot (n=3; mean±s.d; Student’s t-test). (C) Immunoblot analysis with anti-Rab5 antibodies of in vitro-binding assays carried out on post-nuclear supernatants of Jurkat cells using IFT20-GST, ΔCC IFT20-GST and CC IFT20-GST fusion proteins, or GST as negative control. The Ponceau red staining of the same filter is shown to compare the levels of fusion proteins and GST used in the assay. The immunoblot shown is representative of three independent experiments. (D, E) Immunofluorescence analysis of GFP and Rab5 (D) or ATG16L1 and Rab5 (E) in IFT20-GFP and ΔCC IFT20-GFP Jurkat transfectants. Representative medial optical sections and overlay of immunofluorescence and DIC images are shown (IF+DIC). Scale bar: 5 μm. The graphs show the quantification of the weighted colocalization (Mander’s coefficient) of GFP and the early endosome marker Rab5 (D, ≥33 cells/sample, n=3; mean±s.d; Mann–Whitney test), or of ATG16L1 and Rab5 (E, ≥25 cells/sample, n=3; mean±s.d; Student’s t-test) in Jurkat transfectants. **P < 0.01; ***P < 0.001; ****P < 0.0001

To map the interaction of IFT20 with Rab5 we carried out pull-down assays using the IFT20-GST, IFT20-CC-GST and IFT20-ΔCC-GST fusion proteins (Figure 1C). Rab5 interacted both with the GST-tagged isolated CC domain and with the fusion protein lacking the CC domain, indicating that molecular determinants mapping to different parts of IFT20 contribute to its ability to bind Rab5 (Figure 4C). Consistent with this finding, the CC domain of IFT20 was found to be required for its association with early endosomes, as assessed in Jurkat cell transfectants expressing ΔCC IFT20-GFP and stained for Rab5 (Figure 4D). Additionally, ATG16L co-localization with Rab5 was compromised in these cells (Figure 4E). These results indicate that, by interacting with Rab5 and ATG16L1 using both its unstructured portion and its CC domain, IFT20 promotes the association of ATG16L1 with early endosomes.

### IFT20 is required for the interaction of ATG16L1 with autophagy regulators

The autophagy regulator BECLIN 1 assists the localization of ATG16L1 to the sites of autophagosome formation both indirectly and directly. BECLIN 1 forms a complex with VPS34, which induces a local increase in PI(3)P at the isolation membrane (Funderburk et al., 2010), leading to ATG16L1 recruitment through binding to the PI(3)P-binding adaptor WIPI2 (Dooley et al., 2014). Additionally, ATG16L1 interacts with BECLIN 1 in a complex that includes the gap junction protein connexin 43 (Bejarano et al., 2014). Compartment-specific interactors, such as Rab5 for early endosomes in HeLa cells (Otomo et al., 2011; Ravikumar et al., 2008), promote local autophagosome formation by recruiting BECLIN 1.

The finding that IFT20 is required for the association of ATG16L1 with membrane compartments in T cells raises the question of whether IFT20 is implicated in the formation of the autophagy-regulating complexes in which ATG16L1 participates. Since the IFT20-dependent recruitment of ATG16L1 to the Golgi apparatus does not appear to contribute to its pro-autophagic function in T cells (Figure 3G and H), we focused on early endosomes. In particular, we addressed the role of IFT20 in the interaction of ATG16L1 with BECLIN 1 at this membrane compartment. Similar to other cell types (Bejarano et al., 2014), ATG16L1 was found to co-immunoprecipitate and co-localize with BECLIN 1 in T cells. This association was impaired in IFT20KD T cells (Figure 5A and 5B). Conversely, IFT20 deficiency did not affect the ability of ATG16L1 to interact with its downstream autophagy-regulating partner ATG5 (Figure 5C), which is covalently bound to ATG12 (Mizushima et al., 1998). Additionally, neither the interaction of BECLIN 1 with Rab5 (Figure 5D) nor their co-localization (Figure 5E) were affected by IFT20 deficiency. Accordingly, PI(3)P production at early endosomes was comparable to controls in IFT20KD cells, as assessed by quantifying the co-localization of Rab5 with GFP in cells transiently transfected with a GFP-tagged reporter construct encoding the tandem FYVE domains of hepatocyte growth factor-regulated tyrosine kinase substrate, which specifically interacts with PI(3)P (Figure 5F). Together, these results suggest that IFT20 is required for the recruitment of the ATG16L1-ATG5-ATG12 complex to early endosomes that have been tagged for autophagosome formation by Rab5-associated BECLIN 1.

**Figure 5.**
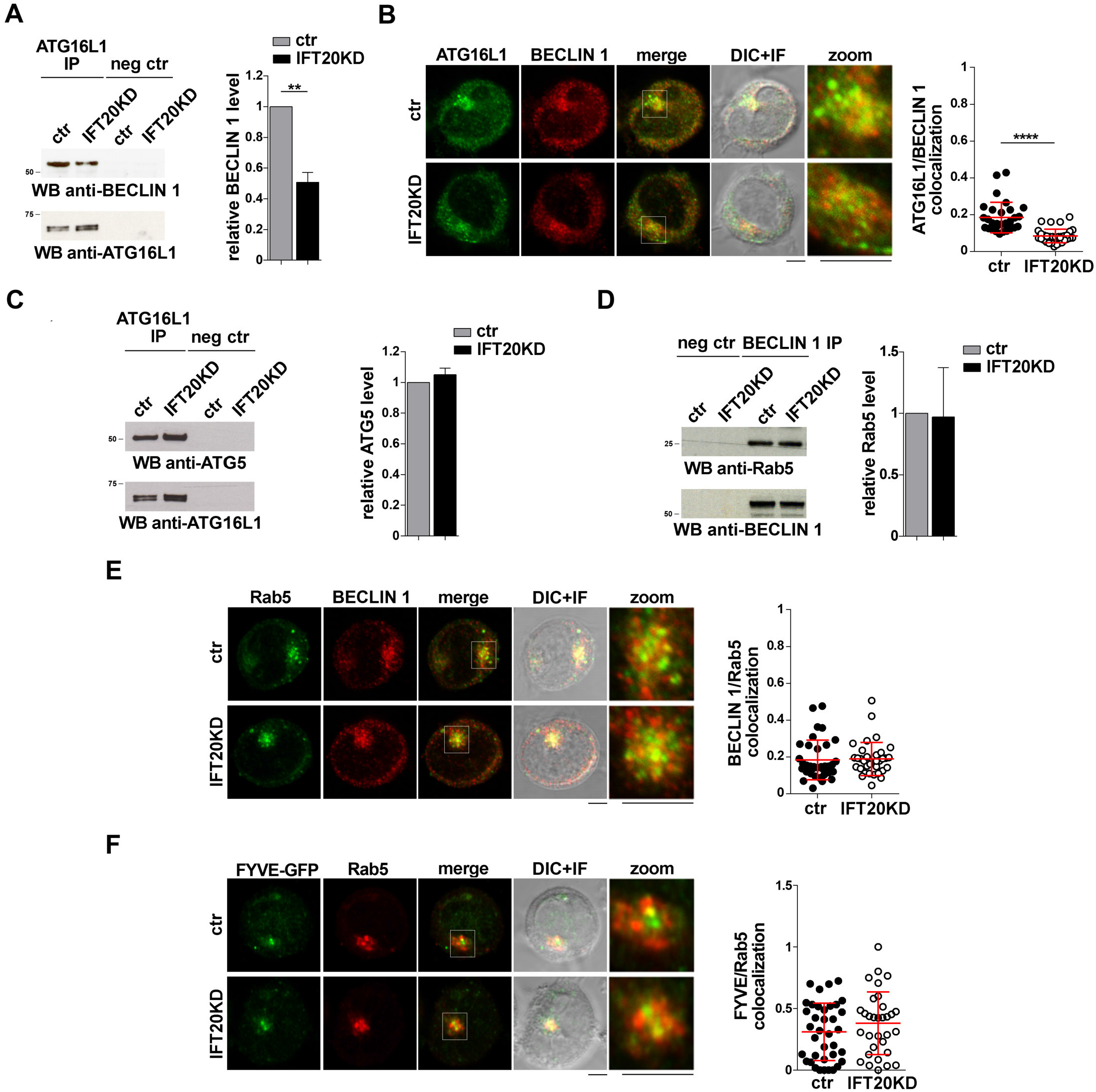
IFT20 recruits ATG16L1 to early endosomes tagged for autophagosome formation. (A) Immunoblot analysis with BECLIN 1 antibodies of ATG16L1-specific immunoprecipitates from lysates of control and IFT20KD Jurkat cells. Preclearing controls are included in each blot (neg ctr). Tested proteins show comparable expression in total cell lysate from ctr and IFT20KD Jurkat cells (Supplementary figure 3A and C). The migration of molecular mass markers is indicated. The quantification of the relative protein expression normalized to ATG16L1 (mean fold ± SD; ctr value=1) is reported for each blot (n=3; mean±s.d; Student’s t-test). (B) Quantification (using Mander’s coefficient) of the weighted colocalization of ATG16L1 with BECLIN 1 in medial confocal sections of control and IFT20KD Jurkat cells (mean±s.d.; ≥30 cells/line; n=3; Mann–Whitney test). Representative images (medial optical sections and overlay DIC+IF) are shown. Scale bar: 5 μm. (C) Immunoblot analysis with ATG5 antibodies of ATG16L1-specific immunoprecipitates from lysates of control and IFT20KD Jurkat cells. Preclearing controls are included in each blot (neg ctr). Tested proteins show comparable expression in total cell lysate from ctr and IFT20KD Jurkat cells (Supplementary figure 3A and D). The migration of molecular mass markers is indicated. The quantification of the relative protein expression normalized to ATG16L1 (mean fold ± SD; ctr value=1) is reported for each blot (n=3; mean±s.d). (D) Immunoblot analysis with Rab5 antibodies of BECLIN 1-specific immunoprecipitates from lysates of control and IFT20KD Jurkat cells. Preclearing controls are included in each blot (neg ctr). Tested proteins show comparable expression in total cell lysate from ctr and IFT20KD Jurkat cells (Supplementary figure 3B and C). The migration of molecular mass markers is indicated. The quantification of the relative protein expression normalized to ATG16L1 (mean fold ± SD; ctr value=1) is reported for each blot (n=3; mean±s.d). (E, F) Immunofluorescence analysis of BECLIN 1 and Rab5 (E) or FYVE-GFP and Rab5 (F) in control and IFT20KD Jurkat cells. Representative medial optical sections and overlay of immunofluorescence and DIC images are shown (IF+DIC). Scale bar: 5 μm. The graphs show the quantification of the weighted colocalization (Mander’s coefficient) of BECLIN 1 (E) or FYVE-GFP (F) and Rab5 in ctr and IFT20KD Jurkat cells(mean±s.d.; ≥32 cells/line; n=3; Mann–Whitney test). ****P < 0.0001

ATG16L1 is involved in determining the site of LC3 lipidation, functioning as a scaffold to recruit the ATG12-ATG5 conjugating enzyme and LC3 to allow for the transfer reaction of LC3 from ATG3 to phosphatidylethanolamine at the phagophore membrane (Fujita et al., 2008; Hanada et al., 2007). The potential impact of IFT20 depletion on the ability of ATG16L1 to interact with LC3 was assessed in co-immunoprecipitation experiments. ATG16L1 was found to co-immunoprecipitate with LC3-I (Figure 6A). This suggests that the pool of ATG16L1 recruited to early endosomes by IFT20 might in turn recruit LC3-I to promote its local cleavage and lipidation to LC3-II. In support of this notion, the interaction of ATG16L1 with LC3-I was significantly impaired in IFT20-deficient T cells (Figure 6A). Consistent with these results, IFT20 deficiency resulted in a decrease in the co-localization of ATG16L1 with LC3 dots (Figure 6B). Importantly, a decrease in the co-localization of LC3 with Rab5 was also observed under these conditions (Figure 6C). Taken together, these results indicate that IFT20 participates in autophagosome formation by targeting ATG16L1 and its partners ATG5-ATG12 to Rab5^+^ and BECLIN-1^+^ vesicles, allowing for the recruitment of LC3 to promote the local generation of isolation membranes at early endosomes.

**Figure 6.**
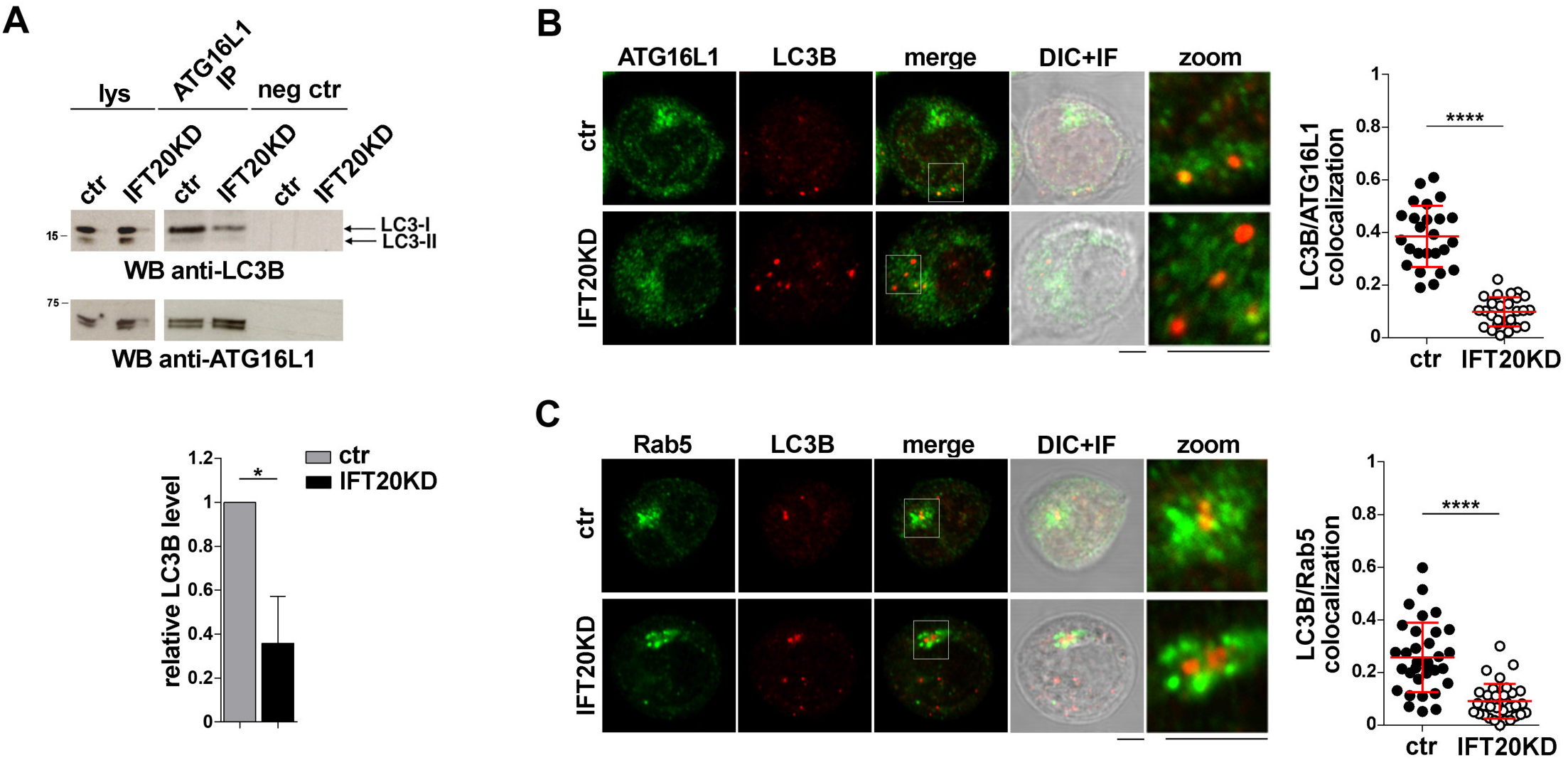
IFT20 is required for LC3 recruitment to ATG16L1 at early endosomes. (A) Western blot analysis with anti-LC3B antibodies of ATG16L1-specific immunoprecipitates from lysates of control and IFT20KD Jurkat cells. Preclearing controls and total cell lysates are included in each blot (neg ctr). The migration of molecular mass markers is indicated. The quantification of the relative protein levels normalized to ATG16L1 (mean fold ± SD; ctr value=1) is shown (n=3; mean±s.d; Student’s t-test). (B,C) Immunofluorescence analysis of ATG16L1 (B) or Rab5 (C) and LC3B in control and IFT20KD Jurkat cells. The graph shows the quantification using Mander’s coefficient of the weighted colocalization of LC3B and ATG16L1 (B) or Rab5 (C) in ctr and IFT20KD Jurkat cells reported as mean value/cell calculated on single dot LC3^+^ (≥25 cells/sample, n=3; mean±s.d; Mann–Whitney test). Representative images (medial optical sections and the overlay DIC+IF) are shown. Scale bar: 5 μm. *P < 0.05; ****P < 0.0001.

## DISCUSSION

We have previously reported that IFT20 indirectly promotes T cell autophagy by regulating the mannose-6-P receptor-dependent transport of acid hydrolases to lysosomes (Finetti et al., 2020). Based on the implication of IFT20 in autophagosome formation in ciliated cells (Pampliega et al., 2013), here we investigated the potential direct implication of IFT20 in this process in the non-ciliated T cell. We show that, under basal conditions, IFT20 interacts with the autophagy regulator ATG16L1 and recruits it to both the Golgi apparatus and early endosomes. For this function IFT20 exploits its CC domain to bind the golgin GMAP210 at the Golgi, and molecular determinants mapping to both the CC domain and the unstructured domain to bind the small GTPase Rab5 at early endosomes. We provide evidence that the IFT20-dependent ATG16L1 localization to early endosomes, but not to the Golgi apparatus, results in the local formation of autophagosomes downstream of BECLIN 1 recruitment and PI(3)P production. The results highlight a dual role of IFT20 in basal T cell autophagy.

IFT20 has been previously demonstrated to regulate both basal and starvation-induced autophagy in ciliated cells (Pampliega et al., 2013). Our data, implicating IFT20 in basal T cell autophagy, underscore the notion that the ciliogenesis machinery is exploited by the non-ciliated T cell beyond immune synapse assembly. In support of this notion, we have recently demonstrated that IFT20 participates in lysosome biogenesis, a function that we found to be shared by ciliated cells (Finetti et al., 2020). The autophagy-related mechanism appears however to differ, at least in part. In ciliated cells the IFT system is required for the transport of autophagy regulators - including VPS34, ATG14, ATG16L1 and ATG7 - to the base of the cilium (Pampliega et al., 2013). Additionally, IFT20 co-localizes with ATG16L1 at small Golgi-proximal vesicles that undergo trafficking to the cilium upon cell starvation, independently of other components of the IFT system (Pampliega et al., 2013). Here we show that IFT20 co-localizes with ATG16L1 at the Golgi apparatus. Additionally, the IFT20-dependent Golgi localization of ATG16L1 does not appear relevant to the pro-autophagic function of IFT20, at least under basal conditions. Indeed, untethering IFT20 from the Golgi through depletion of its interactor GMAP210 did not compromize autophagy, despite the dissipation of ATG16L1 away from the Golgi. The Golgi apparatus has been identified as one of the membrane sources for phagophore elongation in other cell types, with ATG16L promoting the local recruitment and lipidation of LC3 (Davis et al., 2017; Fujita et al., 2008). Our data suggest that other ATG16L pools, recruited to the Golgi apparatus by alternative interactors, such as Rab33 (Itoh et al., 2008), might contribute to the local formation of autophagosomes. However, we have detected minimal co-localization of BECLIN 1 with the Golgi in T cells. As an alternative possibility, Golgi-associated ATG16L may play autophagy-independent functions in these cells, as recently reported in *Dictyostelium discoideum* (Karow et al., 2020).

We found that, as opposed to the Golgi, early endosomes are sites of autophagosome formation in T cells, as shown by the local accumulation of BECLIN 1, PI(3)P, ATG16L1 and LC3. Interestingly, while IFT20 appears dispensable for the association of a functional class III PI3-K complex with early endosomes, it is essential for the recruitment of ATG16L to this location. PI(3)P is known to promote the WIPI2-dependent association of ATG16L with sites of autophagosome formation (Dooley et al., 2014). However, additional interactors participate in its targeting to specific membrane compartments, including Rab33 at the Golgi apparatus, SNX18/Rab11 at recycling endosomes and clathrin at the plasma membrane (Itoh et al., 2008; Knævelsrud et al., 2013; Puri et al., 2013; Ravikumar et al., 2010). Here we identify Rab5 as an early endosome-specific binding partner of ATG16L in T cells, which may account for the association of ATG16L with early endosomes reported for other cell types (Fraser et al., 2019). Of note, Rab5 participates in a complex with BECLIN 1 and VPS34 (Ravikumar et al., 2008). Our results implicate Rab5 not only in marking early endosomes as sites of autophagosome formation by promoting the VPS34-dependent production of PI(3)P, but also in assisting the subsequent recruitment and/or stabilization of ATG16L at these sites using IFT20 as an adaptor.

Our data provide evidence that the association of IFT20 with both the Golgi apparatus and early endosomes is mediated by compartment-specific interactors, namely GMAP210, as previously reported (Follit et al., 2008) and Rab5, respectively. Both these interactions map to the CC domain of IFT20, as shown both in pull-down assays and by co-localization analysis of cells expressing an IFT20 deletion mutant lacking the CC domain. The site of interaction of IFT20 on GMAP210 has been mapped to the CC domain of the latter (Follit et al., 2008; Zucchetti et al., 2019). Hence GMAP210 and IFT20 heterodimerize through their respective CC domain, consistent with the function of CC domains in establishing protein-protein interactions (Mason and Arndt, 2004). IFT20 exploits its CC domain to also interact with Rab5, however the pull-down assays with the GST-tagged IFT20 construct lacking the CC domain show that the unstructured domain also binds Rab5, indicating the cooperation of multiple molecular determinants of IFT20 in this interaction. Of note, at variance with its compartment-specific binding partners, the interaction of IFT20 with ATG16L1 is CC domain-independent and maps to the unstructured N-terminal portion of the protein. While the only protein-protein interaction domain of IFT20 is its CC domain, which is used for binding IFT54 and IFT57 within the IFT complex (Baker et al., 2003; Omori et al., 2008), IFT20 is able to recognize and bind a number of proteins at the Golgi apparatus for sorting and targeting to the ciliary membrane, as exemplified by rhodopsin, opsin and polycystin-2 (Follit et al., 2006; Keady et al., 2011). Hence, similar to these interactors, ATG16L1 might bind directly or indirectly to a molecular determinant within the IFT20 N-terminus that remains to be identified.

Longevity is a key feature of both naive and memory T cells as it ensures the maintenance of the immune repertoire as well as long-lasting protection from pathogens towards which the organism has mounted an effective immune response. Basal autophagy is one of the main mechanisms that underlie this extended T cell survival (Kovacs et al., 2012; Pua et al., 2007; Xu et al., 2014). Our finding of a dual role of IFT20 in autophagy, i.e. promoting autophagosome formation by recruiting ATG16L1 at early endosomes, and allowing for the degradation of the autolysosome contents by controlling lysosome biogenesis (Finetti et al., 2020), highlights IFT20 as a key player in T cell homeostasis.

## Conflict of Interest

The authors declare that the research was conducted in the absence of any commercial or financial relationships that could be construed as a potential conflict of interest.

## Author Contributions

All authors listed have made a substantial, direct and intellectual contribution to the work, and approved it for publication. FF and CB wrote the manuscript. FF prepared the figures.

## Funding

This work was carried out with the support of Fondazione Telethon, Italy (Grant GGP16003) and Associazione Italiana per la Ricerca sul Cancro (Grant IG 20148) to CTB. VC was supported by the Lundbeck Foundation (R209–2015–3505) and the KBVU from the Danish Cancer Society (R146-A9471), and is currently supported by the Fondazione Umberto Veronesi. The support of the “Associazione Italiana per la Ricerca sul Cancro” (AIRC IG-23543) to Francesco Cecconi is also acknowledged.

## Acknowledgements

The authors wish to thank Greg Pazour, Marino Zerial and Antonella De Matteis for generously providing key reagents.

## Notes

### Competing Interest Statement

The authors have declared no competing interest.

