## Supplemental figures and tables for "The intraflagellar transport protein IFT20 recruits ATG16L1 to early endosomes to promote autophagosome formation in T cells"

### SUPPLEMENTARY FIGURE LEGENDS

**Supplementary figure 1.** (A) Representative IFT20 immunoblot on lysates from Jurkat cells, transfected with either empty vector (GFP), with the IFT20-GFP construct (IFT20-GFP), or with the plasmid encoding for ΔCC IFT20-GFP (ΔCC IFT20-GFP). Actin was used as a loading control. (B) Immunoblot analysis of IFT20 in lysates of control and IFT20KD Jurkat cells (mean fold ± SD; one-sample t test; $n > 3$). (C) Representative IFT20 immunoblot on lysates from control and IFT20 knocked-down (KD) cells transiently transfected with empty vector (GFP), with the IFT20-GFP construct or with ΔCC IFT20-GFP vector. Actin was used as a loading control. (D) Immunoblot analysis of GMAP210 in lysates of control and GMAP210KD Jurkat cells (mean fold ± SD; one-sample t test; n=3). The migration of molecular mass markers is indicated; #, non-specific signal. *$P < 0.05$; ***$P<0.0001$

**Supplementary figure 2.** Quantification of Mander's colocalization coefficient (mean ± SD; ≥ 28 cells, n = 3) between BECLIN 1 and GM130 in medial confocal sections of control or IFT20KD Jurkat cells stained with the respective antibodies. Representative images are shown. Size bar: 5 μm.

**Supplementary figure 3.** Representative immunoblot anti-ATG16L1 (A), anti-Rab5 (B), anti-BECLIN 1 (C) and anti-ATG5 (D) on lysates from control and IFT20KD Jurkat cells. The quantification of the relative protein expression is normalized to control (mean fold ± SD; n=3). The migration of molecular mass markers is indicated.

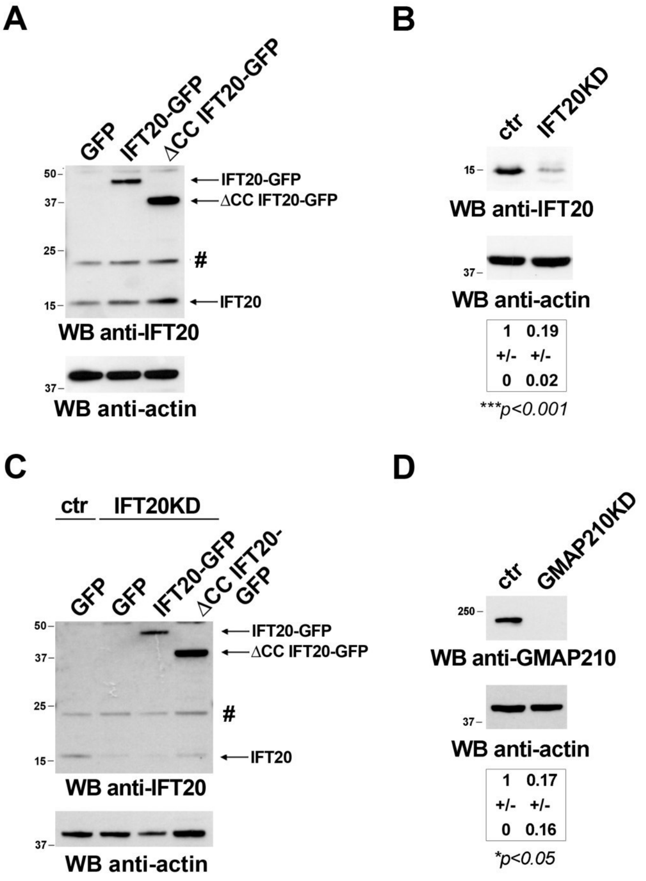


Supplementary figure 1

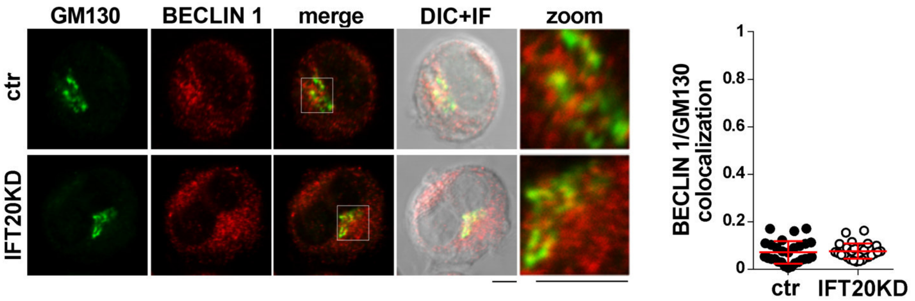


Supplementary figure 2

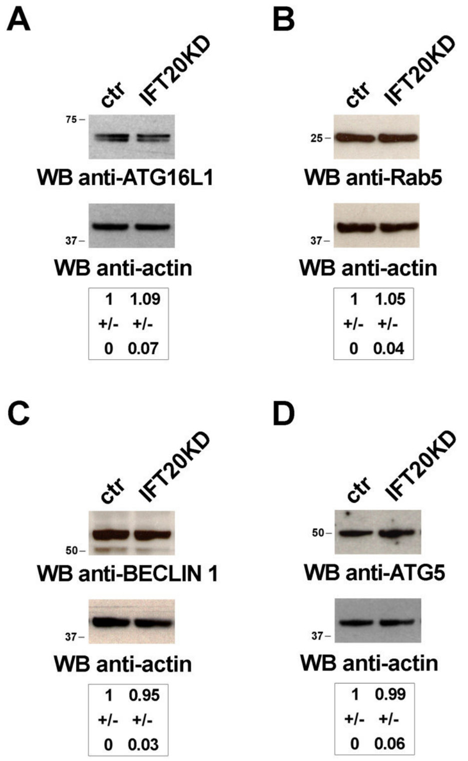


Supplementary figure 3

### SUPPLEMENTARY TABLES

#### Table S1. List of the primers used in this study

| Primers | Sequence |
|---|---|
| **∆CC mutant-GFP XhoI Fw** | CCGCTCGAGATGGCCAAGGACATCCTG |
| **∆CC mutant-GFP KpnI Rev** | CGGGGTACCCCGATGGCCTTCATCTTT |
| **CC mutant-GFP XhoI Fw** | CCGCTCGAGATGGGTGCTCGGAACT |
| **CC mutant-GFP KpnI Rev** | CGGGGTACCCCTTTCTGAAAAATAAATTGGTC |
| **GST-∆CC mutant EcoRI Fw** | CCGGAATTCCGATGGCCAAGGACAT |
| **GST-∆CC mutant XhoI Rev** | CCGCTCGAGTCAGATGGCCTTCATCTTTTC |
| **GST-CC mutant EcoRI Fw** | CCGGAATTCCGATGGGTGCTCGGAA |
| **GST-CC mutant XhoI Rev** | CCGCTCGAGTCATTTCTGAAAAATAAATTGGTCAA |
| **GST-IFT20 EcoRI Fw** | CCGGAATTCCGATGGCCAAGGACAT |
| **GST-IFT20 XhoI Rev** | CCGCTCGAGTCATTTCTGAAAAATAAATTGGTCAA |

**Table S2. List of the antibodies used in this study**

| Antibody | Host Species | Catalogue number | Source | Dilution WB | Dilution IF |
|---|---|---|---|---|---|
| Anti-actin | mouse | MAB1501 | EMD Millipore | 1:10000 | - |
| Anti-ATG5 | mouse | sc-133158 | Santa Cruz | 1:500 | - |
| Anti-Atg16L1 | mouse | sc-393274 | Santa Cruz | 1:500 | 1:50 |
| Anti-Atg16L1 | rabbit | 8089S | Cell Signaling | 1:500 | - |
| Anti-BECLIN 1 | mouse | sc-48341 | Santa Cruz | 1:500 | - |
| Anti-BECLIN 1 | rabbit | 3495T | Cell Signaling | 1:500 | 1:50 |
| Anti-ERK2 | rabbit | sc-154 | Santa Cruz | 1:500 | - |
| Anti-γ-tubulin | mouse | T6557 | Sigma Aldrich | - | 1:200 |
| Anti-giantin | rabbit | ab8084 | Abcam | - | 1:200 |
| Anti-GFP | rabbit | A11122 | Life Technologies | 1:1000 | 1:200 |
| Anti-GMAP210 | mouse | 611712 | BD Biosciences | 1:250 | - |
| Anti-GM130 | rabbit | 610822 | BD Biosciences | 1:500 | 1:100 |
| Anti-IFT20 | rabbit | - | GJ Pazour* | 1:500 | 1:200 |
| Anti-LC3B | rabbit | 3868 | Cell Signaling | 1:500 | 1:200 |
| Anti-Rab5 | mouse | 610724 | BD | 1:2000 | 1:50 |
| Anti-Rab5 | rabbit | 3547S | Cell Signaling | - | 1:200 |

*Program in Molecular Medicine, University of Massachusetts Medical School, Worcester, MA 01605, USA.
